# Plant genotype-specific modulation of *Clonostachys rosea*-mediated biocontrol of septoria tritici blotch disease on wheat

**DOI:** 10.1101/2024.05.28.596162

**Authors:** Sidhant Chaudhary, Mustafa Zakieh, Mukesh Dubey, Dan Funck Jensen, Laura Grenville-Briggs, Aakash Chawade, Magnus Karlsson

**Affiliations:** Department of Forest Mycology and Plant Pathology, Swedish University of Agricultural Sciences, SE-75007, Uppsala, Sweden; Department of Plant Breeding, Swedish University of Agricultural Sciences, SE-23422, Lomma, Sweden; Department of Plant Protection Biology, Swedish University of Agricultural Sciences, SE-23422, Lomma, Sweden

**Keywords:** Biological control agent, *Clonostachys rosea*, genome-wide association study, integrated pest management, septoria tritici blotch, *Zymoseptoria tritici*

## Abstract

**Background:** Beneficial microorganisms can act as biological control agents (BCAs) by directly targeting pathogens or indirectly by enhancing the plant’s defense mechanisms against pathogens. However, efficiencies with which plants benefit from BCAs vary, potentially because of genetic variation in plants for plant-BCA compatibility. The aim of this study was to explore the genetic variation in winter wheat for modulation of *Clonostachys rosea*-mediated biocontrol of septoria tritici blotch disease caused by the fungal pathogen *Zymoseptoria tritici*.

**Results:** In total, 202 winter wheat genotypes, including landraces and old cultivars grown from 1900 onwards in the Scandinavian countries, were tested under greenhouse-controlled conditions. Foliar spray applications of the pathogen and the fungal BCA in two treatments, i.e., *Z. tritici* (Zt) alone and *Z. tritici* along with *C. rosea* (ZtCr) were used to assess the disease progress over time. The absence and presence of *C. rosea* in Zt and ZtCr, respectively, allowed the dissection of variation for plant disease resistance and biocontrol efficacy. The study showed significant phenotypic variation among plant genotypes for disease progression in both Zt and ZtCr treatments. Moreover, disease progress for individual plant genotypes differed significantly between the two treatments, indicating a plant genotype-dependent variation in biocontrol efficacy. For the phenotypic variation in disease progress and biocontrol efficacy, a genome-wide association study using a 20K single-nucleotide polymorphism (SNP) marker array was also performed. In total, five distinct SNP markers associated with disease resistance and four SNP markers associated with *C. rosea* biocontrol efficacy were identified.

**Conclusions:** This work serves as a foundation to further characterize the genetic basis of plant-BCA interactions, facilitating opportunities for concurrent breeding for disease resistance and biocontrol efficacy.

## 1 Background

Septoria tritici blotch (STB) caused by the fungal pathogen *Zymoseptoria tritici* is one of the major fungal foliar diseases of wheat worldwide, which can cause up to 50 % yield losses during severe epidemics in Europe (Fones & Gurr 2015; Savary *et al*. 2019). *Zymoseptoria tritici* goes through several cycles of sexual and asexual reproduction during a growing season, resulting in repeated infection of new plants (Karisto *et al*. 2018). The fungus produces pseudothecia fruiting bodies that release airborne, sexually produced ascospores, while asexual fruiting bodies called pycnidia generate conidia that are dispersed mainly through rain splash (McDonald & Mundt 2016; Karisto *et al*. 2018). Currently, the main control measures include cultivation of disease-resistant wheat varieties and fungicide applications. However, because of its high evolutionary potential, *Z. tritici* can rapidly overcome plant resistance and adapt to single-target fungicides (Torriani *et al*. 2015; Hellin *et al*. 2021; Klink *et al*. 2021). Therefore, new control measures are needed to complement the existing strategies in an integrated pest management (IPM) context.

One such potential measure is the use of microorganisms for biological disease control of STB. Biological control or biocontrol is defined as the exploitation of living organisms to combat pests and pathogens, directly or indirectly, to provide human benefits (Stenberg *et al*. 2021). Biological control can be further subdivided into natural and conservation biocontrol where the resident natural enemies are used to control pathogens, and classical and augmentative biocontrol where mass-reared BCAs are released into target areas (Stenberg *et al*. 2021). Biocontrol of plant diseases is an attractive alternative to chemical control in conventional agriculture and it can also be utilized in organic agricultural practices. In Europe, there are strong political incentives to develop biological control as an important component for sustainable plant production within IPM strategies. For example, the European Green Deal states that the use of synthetic chemical pesticides should be reduced by 50 % by 2030, and biological control is specifically mentioned by the European Commission (2022) in its proposal for a new regulation on sustainable use of plant protection products.

One such BCA is *Clonostachys rosea*, which is an ascomycete fungus with a generalist lifestyle including saprotrophism, plant endophytism and mycoparasitism (Schroers *et al*. 1999; Jensen *et al*. 2021). Certain strains of *C. rosea* can control plant diseases and are currently used for augmentative biological control (Jensen *et al*. 2021). Until now, *C. rosea* has been reported to exhibit biocontrol properties against more than 30 common fungal and oomycete plant pathogens, including *Pythium tracheiphilum* (Møller *et al*. 2003), *Alternaria* spp. (Koch *et al*. 2010), *Botrytis cinerea* (Peng *et al*. 1992), *Fusarium* spp. (Xue *et al*. 2009) and *Bipolaris sorokiniana* (Jensen *et al*. 2016a), on a range of crops including fruits, vegetables, pulses, cereals, oil crops and forest trees (Jensen *et al*. 2021). More recently, *C. rosea* strain IK726 was also shown to control STB on wheat under field conditions (Jensen *et al*. 2019). Mechanistically, *C. rosea* employs different strategies during microbial interactions such as competition for space and nutrients (Sutton *et al*. 1997), antibiosis (Han *et al*. 2020; Saraiva *et al*. 2020), induction of plant defense responses (Wang *et al*. 2019; Kamou *et al*. 2020) and direct parasitism (Barnett & Lilly 1962; Jensen *et al*. 2021).

Most work on augmentative biological control of plant diseases typically involves single or very few plant genotypes. For this reason, it is difficult to explore what effects plant genotypic differences have on the efficacy of BCAs and how this can be exploited in plant breeding. Nevertheless, the limited examples that exist have shown that host plant genotypes can play important roles in the outcome of biocontrol interactions by affecting plant colonization by BCA and pathogen, plant anatomy and physiology and induction of plant defense immune responses (Jensen *et al*. 2016b). Moraga-Suazo *et al*. (2016) reported a differential response towards *C. rosea*-mediated biocontrol of the pitch canker pathogen *F. circinatum* between *Pinus radiata* genotypes. This study also showed that the ability to activate induced systemic resistance (ISR) differed between the pine genotypes, providing indications of the underlying mechanism of the phenomenon. Similarly, Tucci *et al*. (2011) observed differences between five tomato genotypes for enhanced ISR against the grey mold pathogen *B. cinerea* using *Trichoderma atroviride* and *Trichoderma harzianum.* Plant genotype differences for *Trichoderma*-mediated growth promotion in the absence of pathogens are also reported for sugar beet (Schmidt *et al*. 2020) and lentils (Prashar & Vandenberg 2017). From these examples, it is evident that plant genotypes influence the compatibility between plants and beneficial microorganisms. Hence, it is important to consider plant genetic variation for efficient deployment of BCAs.

Breeding for plant crop varieties with enhanced compatibility with beneficial microorganisms offers a long-term, stable and sustainable strategy for efficient plant protection within an IPM context. This requires an understanding of the genetic inheritance of the compatibility trait, and identification of genetic markers useful for marker-assisted breeding or genomic selection approaches. Understanding the nature of the relationship between various agronomic traits and disease resistance traits is an important step towards simultaneous breeding with minimum to no penalty on traditional breeding traits. Recent advances in DNA sequencing technology have enabled large-scale genotyping-by-sequencing approaches useful for large and complex genomes such as *Triticum aestivum* (Lukaszewski *et al*. 2014), which can be exploited in genome-wide association studies (GWAS) of traits such as BCA compatibility.

In the current work, we hypothesized that winter wheat genotypes exhibit variation in their ability to benefit from *C. rosea*-mediated biocontrol of STB disease. Furthermore, we hypothesized that the variation in wheat genotypes affecting *C. rosea* efficacy is genetically distinct from STB disease resistance and inherited in different genomic regions. By screening more than 200 different winter wheat genotypes under controlled conditions, we identified a significant effect of plant genotype on the biocontrol efficacy of *C. rosea* in controlling STB disease. We also identified single nucleotide polymorphism (SNP) markers segregating with the differential response of wheat genotypes toward *C. rosea*-mediated biocontrol, distinct from SNP markers segregating with STB disease resistance. In summary, our results indicate the potential for concurrent breeding of winter wheat varieties with high BCA *C. rosea* compatibility and high STB disease resistance.

## 2 Methods

### 2.1 *Clonostachys rosea* formulation production and application

*Clonostachys rosea* strain IK726 initially isolated from barley roots in Denmark (Knudsen *et al*. 1995) was used in the current work. The strain was revived from a 20 % glycerol conidial suspension stored at -80°C and then maintained on potato dextrose agar media (PDA; BD Difco Laboratories, France) at 20°C in dark conditions. For climate chamber bioassays, a formulation of *C. rosea* IK726 was prepared using a sphagnum peat and wheat bran mixture with some modifications from a previously described method (Jensen *et al*. 2000). Briefly, formulations of *C. rosea* IK726 were prepared from growth on a mixture of sphagnum peat, wheat bran, and water (3:5:12 w/w/w). The mixture was autoclaved twice on two successive days for 20 minutes. Forty grams mixture was put in reagent bottles with caps and inoculated with three agar plugs (five mm diameter) of *C. rosea* from PDA plates. The fungus was incubated for 20 days at 20°C with regular shaking of bottles to promote distribution of *C. rosea* spores. At harvest, the mixture was taken out of the bottles under sterile conditions and was air-dried for two days. Colony forming units (cfu) in the mixture were estimated using a dilution series and the mixture was stored in vacuum sealed bags at 4°C until use.

For foliar application of *C. rosea* in climate chamber bioassays, the formulation was adjusted to 1e10^7^ cfu/ml by adding sterile distilled water. The formulation in water suspension was shaken for 30 minutes to release fungal spores, followed by filtration through miracloth (Merck KGaA, Darmstadt, Germany) to remove larger clumps of mycelium or growth substrate. Tween 20 (Sigma-Aldrich Chemie GmbH, Steinheim, Germany) was added to a final concentration of 0.1 % (v/v) in the *C. rosea* suspension solution as a surfactant immediately before spraying.

### 2.2 *Zymoseptoria tritici* preparation and application

*Zymoseptoria tritici* strain Alnarp 1 was used in this study, which was isolated from STB lesions on leaves of winter wheat collected in 2015 in Lomma, Sweden (Odilbekov *et al*. 2018). The strain was revived from a 50 % glycerol conidial suspension stored at -80°C and then maintained by adding ten µl of the spore suspension in the middle of nine cm diameter YMS medium agar plates, which contained four grams of yeast extract (Merck KGaA, Darmstadt, Germany), four grams of malt extract (Duchefa Biochemie, Haarlem, The Netherlands) and four grams of sucrose (VWR International, Leuven, Belgium) per liter of water (Fagundes *et al*. 2020). After one day of growth on petri plates, the growing culture was spread using a glass spreader to the entire plate by adding one ml sterile water. Inoculated petri plates were incubated at room temperature for ten to twelve days.

For foliar application, *Z. tritici* was harvested by adding sterile distilled water in each petri plate and scraping the mycelial surface with a sterile paint brush to release conidia and spores. The suspension was filtered through a single layer of miracloth (Merck KGaA, Darmstadt, Germany). The concentration of the filtrate was determined using a hemacytometer (Hausser Scientific, Horsham, PA) and was adjusted to 1e10^6^ cfu/ml in the final suspension. Tween 20 (Sigma-Aldrich Chemie GmbH, Steinheim, Germany) was added to a final concentration of 0.1 % (v/v) as a surfactant in the suspension before spraying.

### 2.3 Bioassay experiments

#### 2.3.1 Small-scale biocontrol efficacy screening

An initial experiment in a growth chamber with controlled conditions was performed to optimize concentrations of *Z. tritici* and *C. rosea*, to confirm STB disease development and *C. rosea* biocontrol of STB. Four winter wheat genotypes with varying susceptibility to STB were used: Nimbus (susceptible), Kask (susceptible), SW_150428 (resistant) and Festival (resistant). Four wheat seeds were sown per plastic pot (9×9×8 cm) in potting soil and were placed in trays. Plants were propagated at 60 % relative humidity (RH) with the light period (light intensity of 300 μmol/m^2^) of 16 h at 20°C and dark period of 8 h at 16°C. For each genotype, six treatments were used i.e., 1.) control (mock treatment with no *C. rosea* and no *Z. tritici*), 2.) Cr 1e10^8^ (*C. rosea* at 1e10^8^ cfu/ml), 3.) Zt 1e10^6^ (*Z. tritici* at 1e10^6^ cfu/ml), 4.) Cr 1e10^8^ & Zt 1e10^6^ (*C. rosea* at 1e10^8^ cfu/ml and *Z. tritici* at 1e10^6^ cfu/ml), 5.) Cr 1e10^8^ & Zt 5e10^5^ (*C. rosea* at 1e10^8^ cfu/ml and *Z. tritici* at 5e10^5^ cfu/ml) and 6.) Cr 1e10^8^ & Zt 1e10^5^ (*C. rosea* at 1e10^8^ cfu/ml and *Z. tritici* at 1e10^5^ cfu/ml). With these treatments, it was possible to observe the effect of *C. rosea, Z. tritici* and their various combinations. Five biological replicates were used for each treatment in each genotype.

For *C. rosea* application, 20-day old plants were sprayed with *C. rosea* suspension (control treatment and Zt 1e10^6^ were sprayed with water only) until run-off using a handheld sprayer. Inoculated plants were kept under dark conditions with constant >85 % RH. After 24 h, plants were sprayed with *Z. tritici* suspension (control treatment and Cr 1e10^8^ were sprayed with water only) in the same manner as *C. rosea* application. Inoculated plants were kept under dark conditions with constant > 85% RH. After 48 h, the growth chamber was brought back to standard conditions of 60 % RH with the light period (light intensity of 300 μmol/m^2^) of 16 h at 20°C and dark period of 8 h at 16°C. Percentage of necrotic leaf area was used as a proxy for disease and was visually scored from 0 to 100 % with 5 % step interval. Disease scoring was performed on the (marked) 3^rd^ leaf of each plant at 14, 16, 18, 20, 22, 25, 27 and 30 days post inoculation (dpi) with *Z. tritici*. Using these time points of disease scoring, the relative area under the disease progress curve (rAUDPC) was estimated.

#### 2.3.2 Large-scale biocontrol efficacy screening

A total of 202 winter wheat genotypes were used, which comprised of landraces and cultivars from the Scandinavian countries grown between 1900 and 2012 (Supp. Table 1). The seeds were initially obtained from Nordic Genetic Resources Centre, Alnarp, Sweden, and were later multiplied. Plant seeds were placed on moist filter paper in empty petri plates for four days at 4°C in dark conditions and were then transferred to room temperature for three days for germination. Germinated seedlings were transplanted into plastic pots (9×9×8 cm) filled with potting soil, with two seedlings per pot. Plants were then propagated at 24°C with 60 % RH with a minimum light intensity of 300 μmol/m^2^ s for 16 h and a dark period for 8 h.

The experiment consisted of two treatments i.e., *Z. tritici* (Zt) alone and *Z. tritici* with *C. rosea* (ZtCr). In each treatment, three replicates were used. For treatment Zt, two replicates were evaluated in 2019 (Odilbekov *et al*. 2019) and one replicate in 2022, whereas for ZtCr, all three replicates were evaluated in 2022. In each replicate, the pots were arranged in an augmented design using the R package agricolae (de Mendiburu 2021). Eight to nine blocks of test genotypes with four check genotypes were used in each replicate.

Foliar application of *C. rosea* and *Z. tritici* was done in the same way as in the preliminary small-scale experiment. Plants were sprayed until run-off with *C. rosea* suspension at the concentration of 1e10^7^ cfu/ml in the treatment ZtCr (and with water only in the treatment Zt) and were incubated at 90 % RH. After 24 h, plants were sprayed until run-off with *Z. tritici* at the concentration of 1e10^6^ cfu/ml in both Zt and ZtCr treatments and were incubated at 90 % relative humidity for 48 h. To maintain high humidity, plants were also sprinkled with water four to five times per day. Disease was assessed on two fully developed leaves marked at the base using a marker pen before the inoculation. Percentage of leaf necrotic area was used a proxy for disease and was visually scored from 0 to 100 % with 5 % step interval. Disease scoring was done at three time points in 2019 and at four time points in 2022 at three-day intervals and the disease progress over time was summarized by estimating rAUDPC.

### 2.4 Phenotypic data analysis

#### 2.4.1 Small-scale biocontrol efficacy screening

To check for the genotypic differences between treatments, analysis of variance (ANOVA) was performed using a linear mixed model with genotype and treatment interaction. The model is as follows:

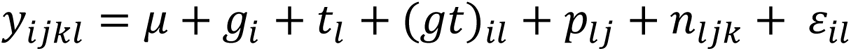

where *y_ijkl_* was the rAUDPC estimate of the *i*-th genotype in the *l*-th treatment, *μ* denotes the overall mean, *g*_*i*_ is the effect of the *i*-th genotype, *t*_*l*_ is the effect of *l*-th treatment, (*gt*)_*il*_ is the interaction effect of of the *i*-th genotype with the *l*-th treatment, *p*_*lj*_ is the effect of the *j*-th pot nested in the *l*-th treatment, *n*_*ljk*_ is the effect of the *n*-th plant nested within the *j*-th pot in the *l*-th treatment, and *ε*_*il*_ is the residual term for which homogenous variance was assumed and was subjected to normal distribution. Pots and plants within pots were considered as random factors in the model. In addition, for multiple comparisons, a post-hoc Tukey’s test among genotypes across treatments was performed.

#### 2.4.2 Large-scale biocontrol efficacy screening

Within each replicate, rAUDPC values were centered and scaled to account for scoring on different days. Therefore, the mean estimate at each replicate level, and ultimately, also at treatment level was centered to 0. However, the absolute rankings of genotypes and the difference among genotypes was still maintained to study genotype level differences. The linear mixed model analysis using Kenward-Roger’s approximation of the degrees of freedom (Kenward & Roger 1997) to estimate best linear unbiased estimators (BLUEs) was performed on these centered and scaled rAUDPC values in the following way:

##### 2.4.2.1 Intra treatment

ANOVA was performed for all rAUDPC separately in each treatment using the following linear mixed model:

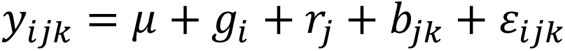

where *y_ijk_* was the phenotypic performance of the *i*-th genotype in the *k*-th block nested within the *j*-th replicate, *μ* denotes the overall mean, *g*_*i*_ is the effect of the *i*-th genotype, *r*_*j*_ is the effect of the *j*-th replicate, *b*_*jk*_ is the effect of the *k*-th block nested within the *j*-th replicate, and *ε*_*ijk*_ is the residual term. Blocks nested within replicates were treated as a random factor. Broad-sense heritability was also estimated in treatments Zt and ZtCr as H^2^_P_ after Piepho & Möhring (2007) and H^2^_c_after Cullis *et al*. (2006).

##### 2.4.2.2 Inter treatment

To check for the genotypic differences between treatments, a full mixed model with genotype and treatment interaction was applied. Genotypes that were not present in both treatments were removed before the analysis (183 genotypes overlapping between treatments). The model is as follows:

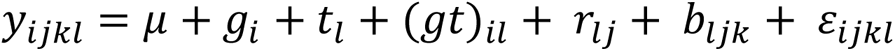

where *y_ijkl_* was the phenotypic performance of the *i*-th genotype in the *k*-th block nested within the *j*-th replicate in the *l*-th treatment, *μ* denotes the overall mean; *g*_*i*_ is the effect of the *i*-th genotype, *t*_*l*_ is the effect of *l*-th treatment, (*gt*)_*il*_ is the interaction effect of of the i-th genotype with the *l*-th treatment, *r*_*lj*_ is the effect of the *j*-th replicate nested in the *l*-th treatment, *b*_*ljk*_ is the effect of the *k*-th block nested within the *j*-th replicate in the *l*-th treatment, and *ε*_*ijkl*_ is the residual term. Blocks nested within replicates in treatments were treated as a random factor. To estimate differences between treatments for each genotype, a post-hoc Tukey’s test was also performed. These inter-treatment genotype contrasts (Zt - ZtCr) were used as estimators for biocontrol efficacy effect.

All statistical analyses were performed using the statistical software R version 4.1.1 “Kick Things” (R Core Team 2021). The linear mixed model analysis was performed using lmer package (Bates *et al*. 2021) and its extension lmertest (Kuznetsova *et al*. 2020). In addition, post-hoc comparisons among genotypes at treatment level and individual genotype comparison between treatments were performed using emmeans (Lenth 2022) and cld (Hothorn *et al*. 2021) R packages. Tidyverse suite (Wickham 2021) was used for most of the data processing and visualization alongside other dependency packages.

### 2.5 Genome-wide association analysis

For GWA mapping, the Genome association and prediction integrated tool (GAPIT) in the R environment was used (Wang & Zhang 2021). The wheat genotypes were previously genotyped using a 20K SNP marker array (Odilbekov *et al*. 2019). SNP markers with > 20 % missing alleles were removed. The remaining missing values were imputed using GAPIT to major allele. For GWA analyses, a threshold of 5 % minor allele frequency was applied. After quality checks, 7360 SNP markers were left for the GWAS (Supp. Table 2). In total, five different models were used: GLM (Price *et al*. 2006), MLM (Yu *et al*. 2006), MLMM (Segura *et al*. 2012), FarmCPU (Liu *et al*. 2016) and BLINK (Huang *et al*. 2019). GLM and MLM are single locus GWA models while MLMM, FarmCPU and BLINK are multiple loci GWA models. The kinship matrix (K) and the first ten principal components (PC) were used as covariates to adjust for familial relatedness and population structure. Only genotypes where SNP marker information was available were included in the GWA analyses. Genotypic marker data was available for 188 genotypes from treatment Zt and 173 individuals from treatment ZtCr and inter-treatment genotype contrasts for biocontrol efficacy. Alongside Bonferroni threshold (0.05/number of SNP markers) for marker-trait association, a less stringent threshold of negative log (1/number of SNP markers) was used to account for relatively low sample size and over stringency of Bonferroni test (Yang *et al*. 2011; Wang *et al*. 2012). For each SNP marker significant at negative log threshold, allelic level comparisons with one-way ANOVA were also made for associated traits.

### 2.6 Identification of candidate genes

We used two methods to define regions in which to search for genes localized at significant marker trait associations. Firstly, as per linkage disequilibrium based criteria explained in Alemu *et al*. (2021), a region of ± 1.6 cM flanking the significantly associated SNP marker was defined as a single quantitative trait locus (QTL) in this germplasm using the same SNP marker chip. The physical positions of SNP markers flanking the ± 1.6 cM region were identified by mapping SNP marker sequences against the *T. aestivum* IWGSC CS RefSeq v2.1 genome (GCF_018294505.1) using the BLAST algorithm. Additionally, a more stringent criteria of ± 100 Kbp was also applied to define regions where the physical positions of flanking SNP markers were identified in the same manner and were used in gene annotation. The genes localized within the physical location of flanking SNP markers were filtered using the gene annotation data of *T. aestivum* (version 55) from the EnsemblPlants database. Only the protein coding gene models assigned to high confidence according to International Wheat Genome Sequencing Consortium (IWGSC) were used for annotation.

The predicted amino acid sequences of the filtered genes were annotated using the BLAST algorithm at the National Centre for Biotechnology Information, and by Conserved Domain Search (Lu *et al*. 2020), the SMART analysis tool (Letunic *et al*. 2021) and InterproScan (Jones *et al*. 2014).

## 3 Results

### 3.1 Biocontrol of septoria tritici blotch disease

To evaluate biocontrol of STB by *C. rosea*, a pilot experiment was performed using two susceptible (Nimbus and Kask) and two resistant genotypes (SW_150428 and Festival). The experiment resulted in STB disease development over time (rAUDPC) in susceptible cultivars and a significant reduction in STB disease in treatments with *C. rosea* (Figure 1, Supp. Table 3). ANOVA test on rAUDPC showed significant genotype-treatment interaction (*P* < 0.001), indicating differences among genotypes and treatments within genotypes, which were further explored using multiple comparisons Tukey test. No disease development in treatments mock control and Cr 1e10^8^ in any of the four genotypes (Figure 1, Supp. Table 3) was detected. In resistant genotypes SW_150428 and Festival, no significant disease development was observed in any of the treatments (Figure 1, Supp. Table 3). The highest rAUDPC value was observed in the treatment Zt 1e10^6^ for susceptible genotypes Nimbus (0.48) and Kask (0.21), which was significantly (*P* < 0.05) higher than the mock-inoculated controls (Figure 1, Supp. Table 3). These rAUDPC estimates in the treatment Zt 1e10^6^ were significantly higher than the three treatments (Cr 1e10^8^ & Zt 1e10^6^, Cr 1e10^8^ & Zt 5e10^5^ and Cr 1e10^8^ & Zt 1e10^5^) that involved *C. rosea*, exhibiting the biocontrol effect of *C. rosea* in controlling STB (Figure 1, Supp. Table 3).

**Figure 1:**
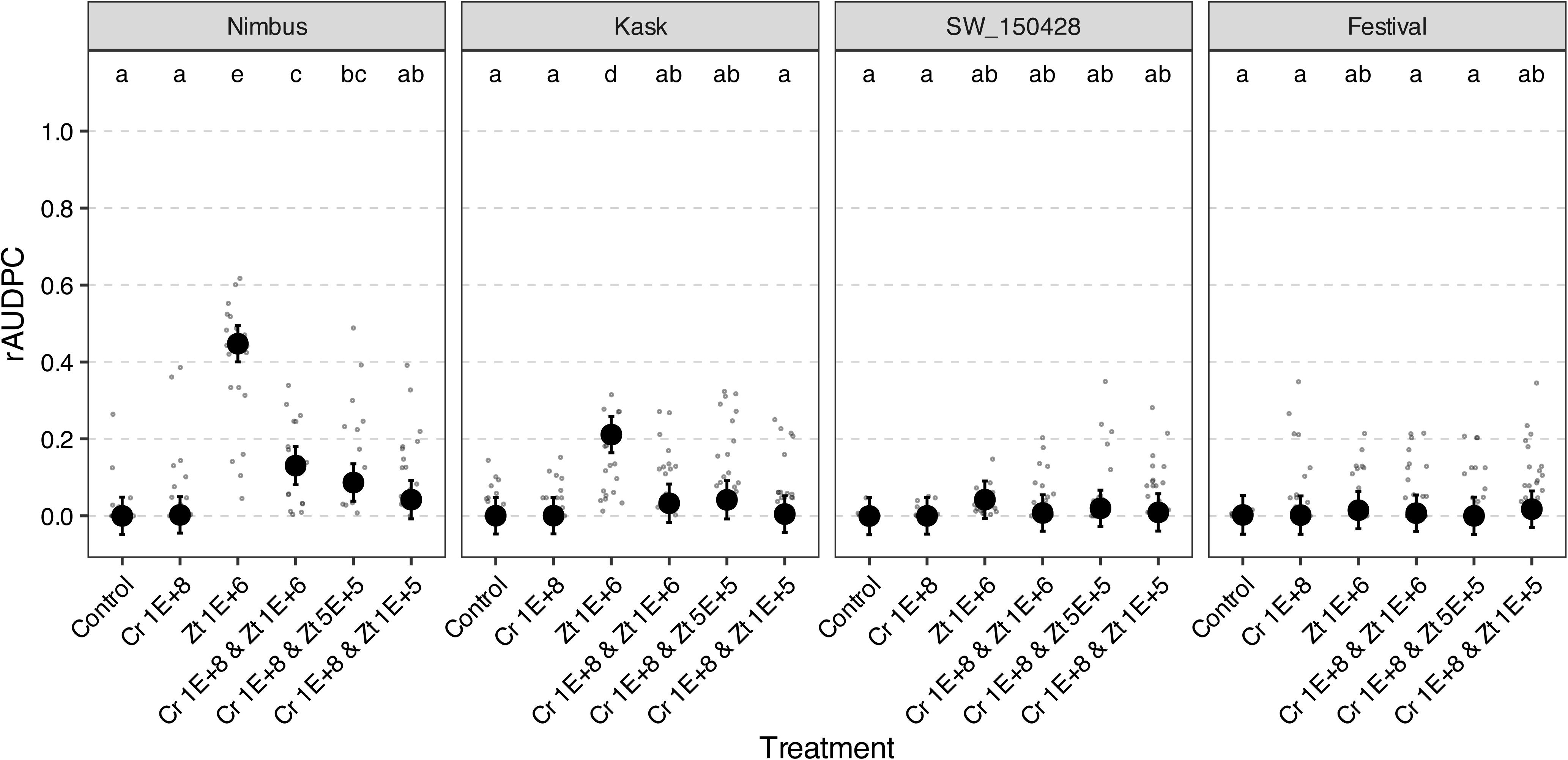
Biological control of septoria tritici blotch disease on wheat. Four winter wheat genotypes with low (Nimbus and Kask) and high (SW_150428 and Festival) disease resistance, respectively, was inoculated with varying concentrations of the biological control agent *C. rosea* and *Z. tritici*. Relative area under disease progress curve (rAUDPC) was used as an estimate of disease severity. Grey points represent raw estimates of rAUDPC values from technical replicates. Black points represent the model estimated means and error bars represent 95 % confidence intervals. Treatments not sharing the same letters indicate significant difference (*P* < 0.05) across genotypes as determined by Tukey’s post-hoc pairwise comparisons test.

### 3.2 Variation among wheat genotypes for septoria tritici blotch disease and *C. rosea* biocontrol efficacy

Application of *Z. tritici* alone (treatment Zt) to 202 winter wheat genotypes resulted in disease development with varying degree among genotypes. Evaluation of rAUDPC in treatment Zt using linear mixed model showed significant variation (*P* < 0.001) for disease severity among genotypes (Table 1, Supp. Figure 1A, Supp. Table 4). rAUDPC in treatment Zt showed moderate to high heritability (H^2^_p_= 0.67, H^2^_c_= 0.59) and these results were in significant strong positive correlation (*R* = 0.69, *P* < 0.001) with STB rAUDPC data reported in Odilbekov et al. (2019) where the same plant material was used for STB disease assessment (Table 1, Figure 2A).

**Figure 2:**
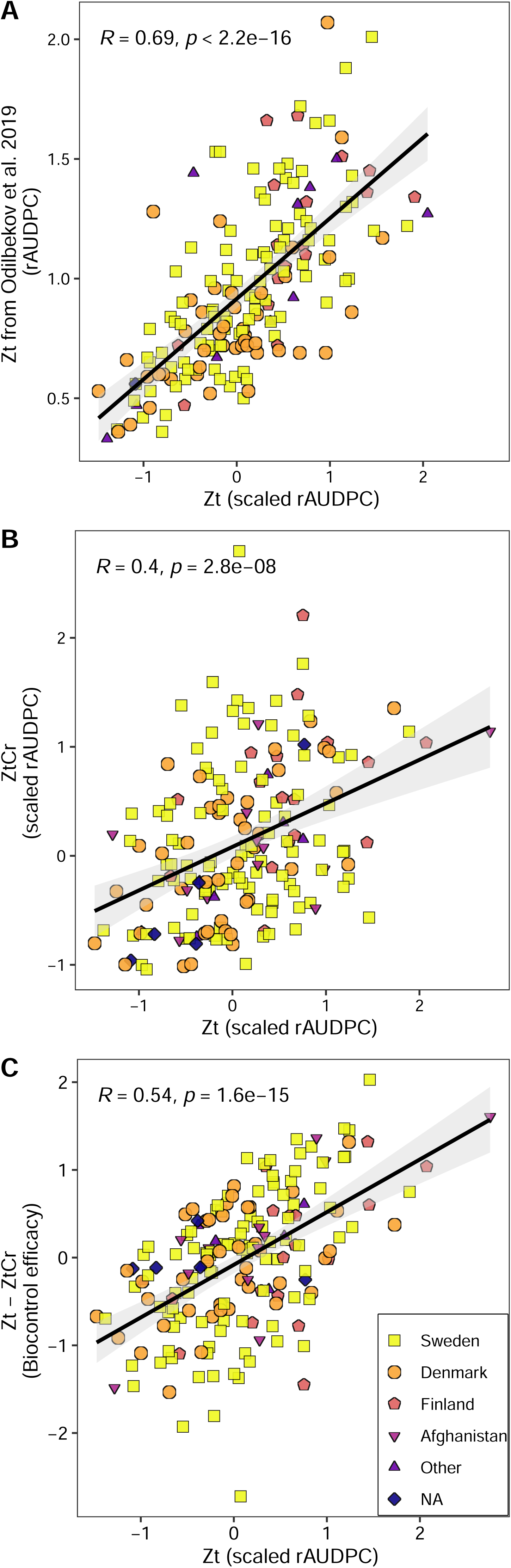
Correlations between treatments and traits. Pearson’s correlation for relative area under disease progress curve (rAUDPC) between different treatments and traits was calculated. (A) Correlation between scaled rAUDPC estimates in treatment Zt (*Z. tritici* only) against rAUDPC estimates in Odilbekov *et al*., (2019), (B) Correlation between scaled rAUDPC estimates in treatment Zt (*Z. tritici* only) against rAUDPC estimate in treatment ZtCr (*Z. tritici* along with *C. rosea*), and (C) Correlation between scaled rAUDPC in treatment Zt (*Z.tritici* only) against biocontrol efficacy estimates (Zt - ZtCr) of *C. rosea* in controlling septoria tritici blotch. Points with different shapes and color represent the country of origin of wheat genotypes.

**Table 1:** Analysis of variance summary from linear mixed models for septoria tritici blotch disease resistance and biocontrol.

| Model<sup>1</sup> | Parameter | Sum of Squares | Mean Squares | NumDF<sup>2</sup> | DenDF<sup>3</sup> | F-value | P-value |
| --- | --- | --- | --- | --- | --- | --- | --- |
| ***Intra treatment*** |  |  |  |  |  |  |  |
| Zt | Rep | 3.4703 | 1.73515 | 2 | 17.13 | 63.1694 | 1.24E-08*** |
|  | Gen | 15.0971 | 0.07511 | 201 | 401.14 | 2.7347 | <2.2E-16*** |
|  | H<sup>2</sup><sub>P</sub> | 0.67 |  |  |  |  |  |
|  | H<sup>2</sup><sub>C</sub> | 0.59 |  |  |  |  |  |
| ZtCr | Rep | 0.7074 | 0.3537 | 2 | 19.93 | 12.7986 | 0.0002662*** |
|  | Gen | 15.3231 | 0.08419 | 182 | 433.97 | 3.0465 | <2.20E-16*** |
|  | H<sup>2</sup><sub>P</sub> | 0.74 |  |  |  |  |  |
|  | H<sup>2</sup><sub>C</sub> | 0.62 |  |  |  |  |  |
| ***Inter treatment*** |  |  |  |  |  |  |  |
|  | Trt (Rep) | 17.6382 | 0.09691 | 182 | 813.33 | 3.4507 | <2.20E-16*** |
|  | Gen | 0.4304 | 0.43041 | 1 | 36.72 | 15.3263 | 0.0003775*** |
|  | Trt | 7.3341 | 0.0403 | 182 | 813.33 | 1.4348 | 0.0005701*** |
|  | Gen x Trt | 3.6963 | 0.92407 | 4 | 36.17 | 32.9048 | 1.35E-11*** |

<sup>1</sup>Models include intra treatment models fit separately in treatments Zt (disease with only *Z. tritici*), ZtCr (disease with *Z. tritici* in presence of biocontrol agent *C. rosea*). Inter treatment model includes Genotype X Treatment interaction.
<sup>2</sup>Numerator degrees of freedom.
<sup>3</sup>Denominator degrees of freedom.
Abbreviations; Rep: Replicate, Gen: Genotype, H<sup>2</sup><sub>P</sub>: Heritability (Piepho & Möhring, 2007), H<sup>2</sup><sub>C</sub>: Heritability (Cullis *et al.*, 2006), Trt: Treatment.

Application of *C. rosea* formulation to the leaves before *Z. tritici* application (treatment ZtCr) also resulted in significant (*P* < 0.001) differences in disease severity between wheat genotypes with moderate to high heritability (H^2^_p_= 0.74, H^2^_C_ = 0.62) (Table 1, Supp. Figure 1B, Supp. Table 5). There was a significant moderate positive correlation (*R* = 0.4, *P* < 0.001) between the treatments Zt and ZtCr (Figure 2B). This moderate positive correlation reflects the changes in disease development in wheat genotypes in presence of *C. rosea*.

### 3.3 Variation among wheat genotypes for *C. rosea* biocontrol efficacy

To quantify differences in rAUDPC values for each genotype between the treatments with *Z. tritici* alone (Zt) and *Z. tritici* with BCA *C. rosea* (ZtCr), a linear mixed model with treatment and genotype interaction effect was employed. A total of 183 genotypes were overlapping between the treatments and were used in the analysis. Identical to the within-treatment analysis in the previous section, significant (*P* < 0.001) variation was observed for STB disease development estimates (rAUDPC) in the treatments Zt and ZtCr. The model also showed significant interaction (*P* < 0.001) between genotype and treatment effect indicating differences in rAUDPC values for genotypes between treatments Zt and ZtCr (Table 1).

rAUDPC differences between treatments (Zt - ZtCr) for each genotype were used as an estimator for biocontrol efficacy and were estimated using Tukey’s multiple comparison test. Post-hoc comparison of 183 wheat genotypes for rAUDPC between Zt and ZtCr revealed a varying degree of disease difference between treatments (i.e. biocontrol efficacy) ranging from more disease in Zt for some genotypes to more disease in ZtCr (negative values) for other genotypes (Figure 3, Supp. Table 6). In particular, seven genotypes (NGB9123, NGB2317, NGB8937, NGB9079, NGB1, NGB17 and NGB6704) had a significant (*P* < 0.05) positive effect of *C. rosea* biocontrol efficacy as they showed higher rAUDPC estimates in treatment Zt than ZtCr (Figure 3, Supp. Table 6). On the other hand, eleven genotypes (NGB6705, NGB6729, NGB6724, NGB15071, NGB9078, NGB23353, NGB348, NGB6699, NGB14118, NGB13445 and NGB8189) had a significant (*P* < 0.05) negative effect of *C. rosea* with higher rAUDPC estimates in treatment ZtCr than Zt (Figure 3, Supp. Table 6). *Clonostachys rosea* biocontrol efficacy estimates were also found to be in significant moderate positive correlation (*R* = 0.54, *P* < 0.001) with rAUDPC estimates from treatment Zt, suggesting that susceptible genotypes benefit more from *C. rosea* application as more reduction in disease was observed (Figure 2C).

**Figure 3:**
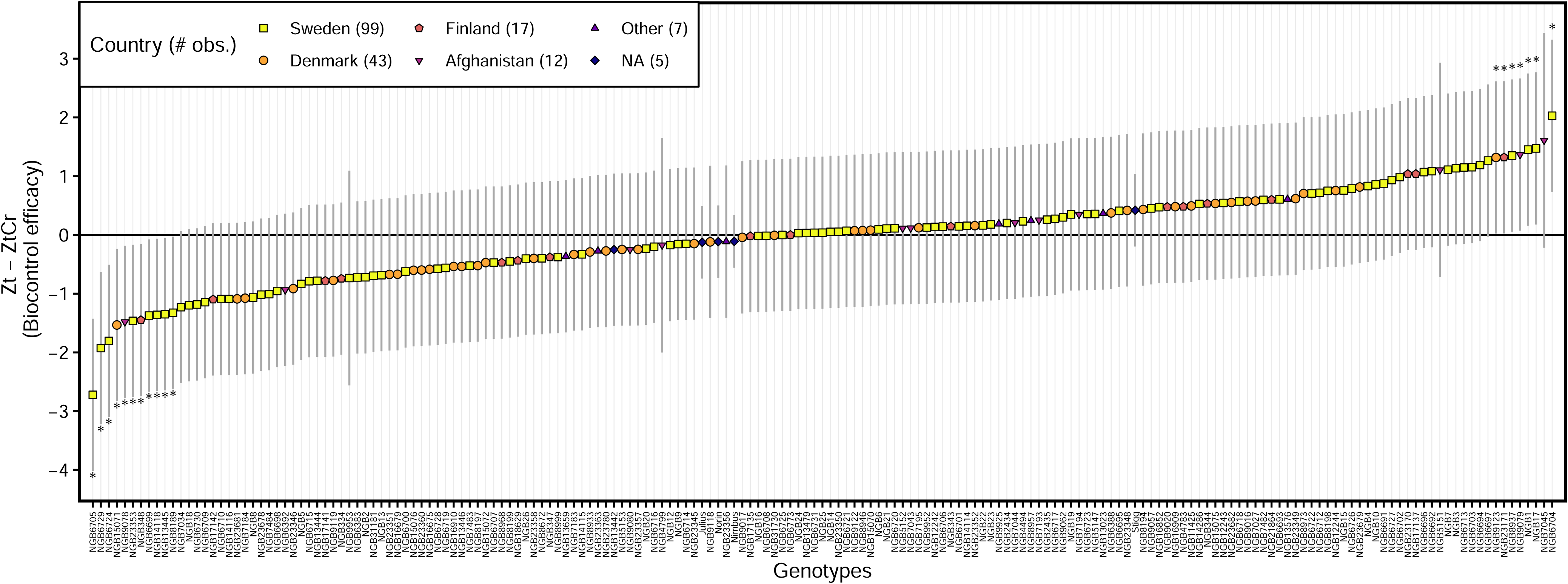
Biocontrol of septoria tritici blotch by *Clonostachys rosea*. Biocontrol efficacy estimates (Zt - ZtCr) of *C. rosea* in controlling septoria tritici blotch for 183 winter wheat genotypes was calculated. Scaled relative area under disease progress curve (rAUDPC) was estimated in treatments Zt (*Z. tritici* alone) and ZtCr (*Z. tritici* along with *C. rosea*) for each wheat genotype using linear mixed models. Post-hoc comparisons using Tukey’s test were used to estimate differences between treatment Zt and ZtCr for each genotype and were used as estimators for biocontrol efficacy. Points represent the model estimated means and error bars represent 95 % confidence intervals for each genotype. Genotypes with ‘*’ (*P* < 0.05) highlight significant difference in rAUDPC estimates between treatments Zt and ZtCr and therefore, represent significant biocontrol efficacy. Points with different shapes and color represent the country of origin of wheat genotypes.

### 3.4 Genome-wide marker trait associations

For phenotypic rAUDPC estimates in treatments with *Z. tritici* alone (Zt), *Z. tritici* along with BCA *C. rosea* (ZtCr) and biocontrol efficacy estimator (Zt - ZtCr), GWA analysis was performed using 20K SNP marker array genotyping data with 7360 markers. The GWAS detected eleven SNP markers (eight of these SNP markers with multi-model GWAS) that were significantly (*P* ≤ 0.00014, after *P* ≤ 1/n, where n = 7360 is the number of SNP markers) associated to rAUDPC variation (Table 2). Seven out of these eleven SNP marker-trait associations were co-detected by more than one GWAS model (Table 2).

**Table 2:**
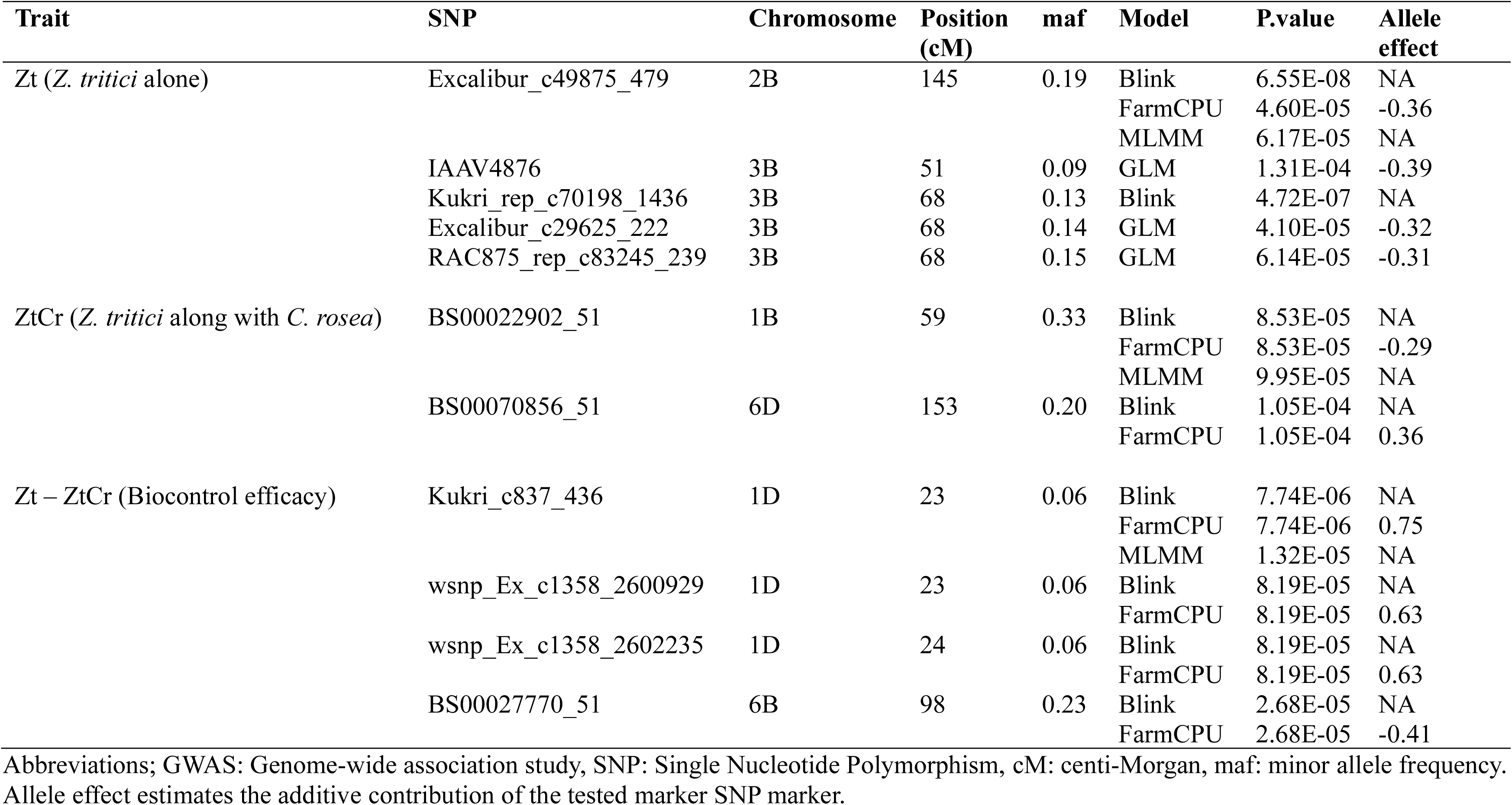
Summary of GWAS results for significant SNP markers associated with Zt (*Z. tritici* alone), ZtCr (*Z. tritici* along with *C. rosea*) and biocontrol efficacy (Zt - ZtCr) estimates in 173 winter wheat genotypes.

For rAUDPC in treatment Zt, five SNP markers at three locations i.e., Excalibur_c49875_479 (chromosome 2B with BLINK, FarmCPU and MLMM model), IAAV4876 (chromosome 3B at 51 cM with GLM model), Excalibur_c29625_222 (chromosome 3B at 68 cM with GLM model), Kukri_rep_c70198_1436 (chromosome 3B at 68 cM with BLINK model) and RAC875_rep_c83245_239 (chromosome 3B at 68 cM with GLM model), were found to be significantly associated (*P* ≤ 0.00014) (Table 2, Figure 4A). At allelic level, genotypes that carried the AA allele for SNP marker IAAV4876 exhibited significantly (*P* < 0.05) less rAUDPC (Supp. Figure 2B).

**Figure 4:**
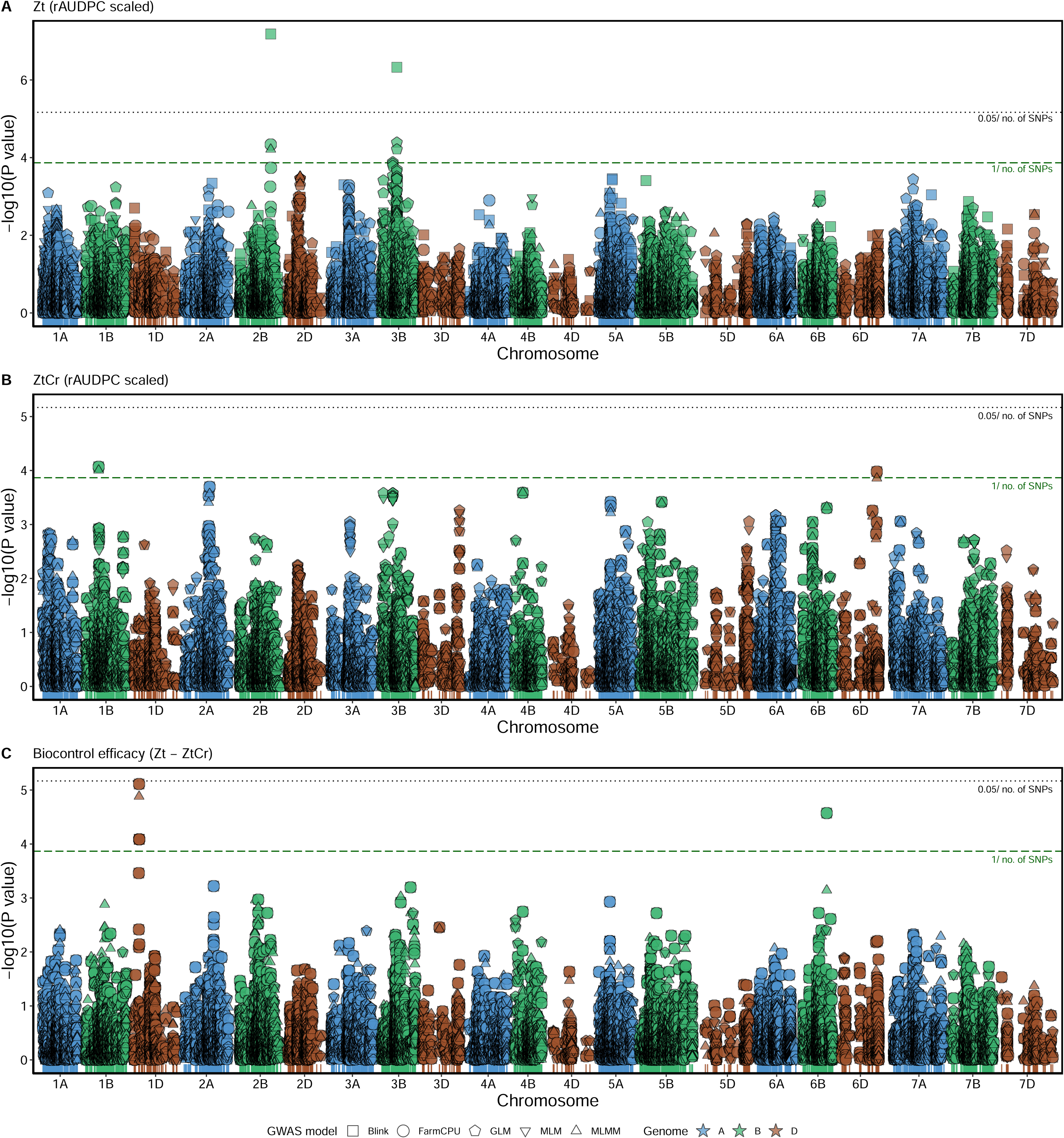
Manhattan plot for marker-trait association between for scaled relative area under disease progress curve (rAUDPC) of wheat genotypes in (A) treatment Zt (*Z. tritici* alone), (B) treatment ZtCr (*Z. tritici* along with *C. rosea*) and (C) Biocontrol efficacy (Zt – ZtCr) and 7360 single nucleotide polymorphism (SNP) markers from the genome– wide association study (GWAS) models. Black dotted line depicts the Bonferroni significance threshold (*P* = 0.05/n, where n = 7360 is the number of SNP markers), green dashed line depicts less stringent threshold (*P* = 0.00014, after *P* = 1/n, where n = 7360 is the number of SNP markers).

For treatment ZtCr in presence of BCA *C. rosea*, two SNP markers i.e., BS00022902_51 (chromosome 1B with BLINK, FarmCPU and MLMM model) and BS00070856_51 (chromosome 6D with BLINK and FarmCPU model), were significantly (*P* ≤ 0.00014) associated with rAUDPC estimates (Table 2, Figure 4B). At allelic level, genotypes that carried the allele TT for SNP marker BS00070856_51 exhibited significantly (*P* < 0.05) less rAUDPC (Supp. Figure 2G).

For biocontrol efficacy (Zt - ZtCr), significant (*P* ≤ 0.00014) SNP marker-trait associations were detected at two locations i.e., at chromosome 1D by SNP markers Kukri_c837_436 (BLINK, FarmCPU and MLMM model), wsnp_Ex_c1358_2600929 (BLINK and FarmCPU) and wsnp_Ex_c1358_2602235 (BLINK and FarmCPU), and at chromosome 6B by SNP marker BS00027770_51 (BLINK and FarmCPU) (Table 2, Figure 4C). At allelic level, genotypes that carried the alleles GG and GT for SNP marker Kukri_c837_436, alleles AA and AG for SNP marker wsnp_Ex_c1358_2600929 and alleles CC and CT for SNP marker wsnp_Ex_c1358_2602235 showed significantly (*P* < 0.05) more biocontrol efficacy (Supp. Figure 2H-J). Moreover, genotypes with allele AA for SNP marker BS00027770_51 showed significantly (*P* < 0.05) more biocontrol efficacy (Supp. Figure 2K).

### 3.5 Gene content in genomic regions with associated SNP markers

The regions around seven locations (three locations for treatment Zt, two locations for treatment ZtCr and two locations for biocontrol efficacy) with significant marker-trait associations were defined using linkage disequilibrium-based criteria defined in Alemu et al. (2021) and a less stringent criteria of ± 100 Kbp flanking the significant markers were used to explore genes. In regions using the criteria of ± 1.6 cM flanking the significant SNP markers, the regions spanned from 0.7 Mbp (for biocontrol efficacy at 1D) to 88.4 Mbp (treatment Zt at chromosome 3B at 68 cM). In total, the number of genes in these regions with assigned high confidence according to IWGSC were 1290 for treatment Zt, 92 for treatment ZtCr and 61 for biocontrol efficacy (Zt – ZtCr) (Supp. Table 7).

Using the more stringent criteria of ± 100 Kbp to define regions, twenty genes were found to be localized within the genomic regions surrounding the physical position of GWAS-identified SNP markers significantly (*P* ≤ 0.00014) associated with treatments Zt, ZtCr and biocontrol efficacy (Zt – ZtCr) (Table 3). Fifteen genes (TraesCS2B02G608100, TraesCS2B02G608200, TraesCS3B02G006500, TraesCS3B02G006600, TraesCS3B02G307000, TraesCS3B02G307100, TraesCS3B02G307200, TraesCS3B02G307300, TraesCS3B02G307400, TraesCS3B02G307500, TraesCS3B02G307600, TraesCS3B02G307700, TraesCS3B02G307800, TraesCS3B02G307900 and TraesCS3B02G308000) were localized within ± 100 Kbp intervals of SNP markers associated with treatment Zt for disease severity (Table 3). TraesCS3B02G307000 was predicted to encode a protein with sequence similarity to a plant homeodomain (PHD) Zinc finger-type pathogenesis-related transcription factor in *Arabidopsis thaliana*. Other predicted proteins with putative functions in plant defense and stress mitigation included a NUDIX domain-containing protein, a cytochrome P450 monooxygenase, a dynamin-like GTPase, VQ motif-containing protein and a carotenoid cleavage dioxygenase (Table 3).

**Table 3.**
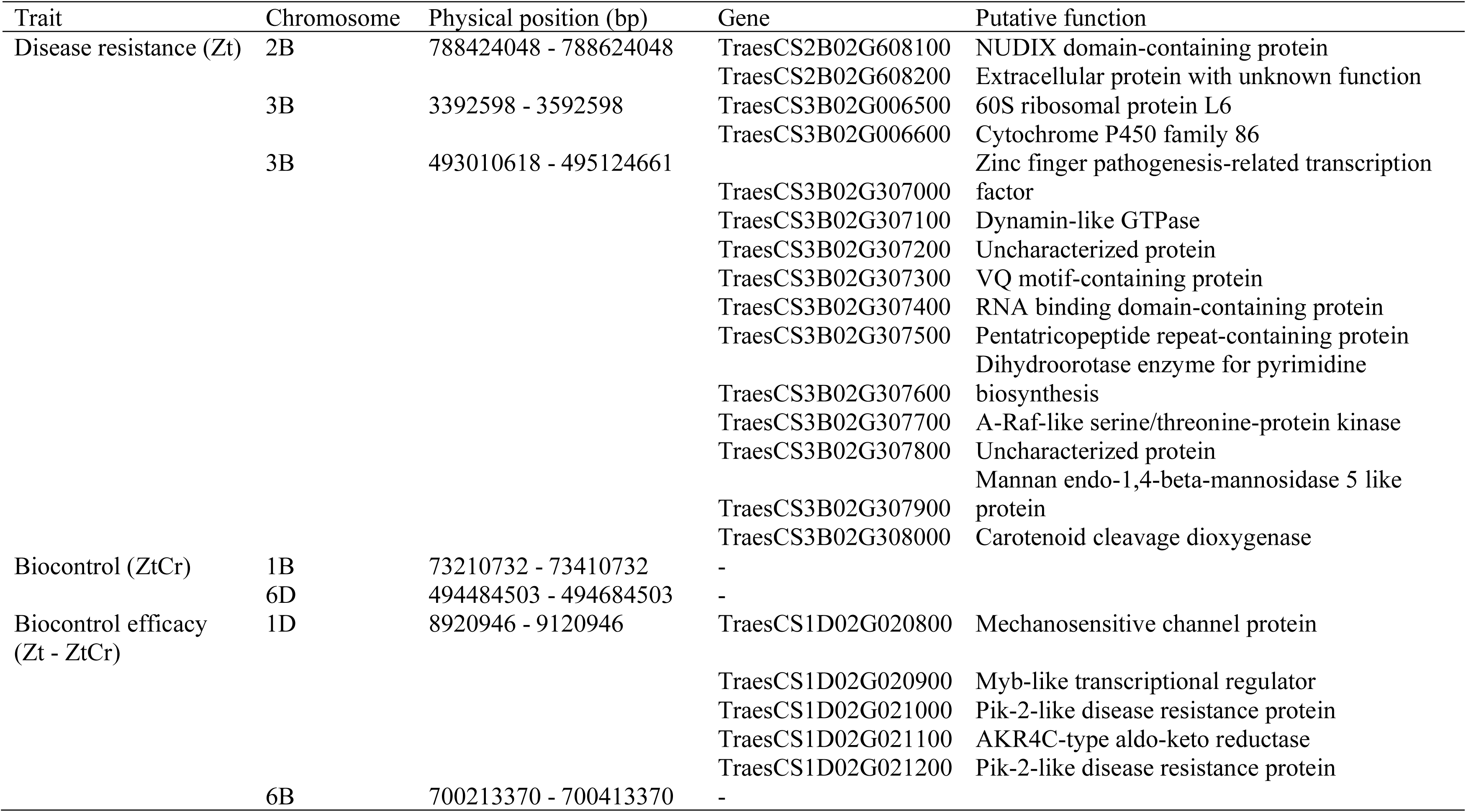
Gene content of wheat genomic regions segregating with septoria tritici blotch disease resistance and biocontrol in ± 100 Kbp interval.

No genes were identified in the genomic regions segregating with biocontrol treatment ZtCr (Table 3). However, five genes (TraesCS1D02G020800, TraesCS1D02G020900, TraesCS1D02G021000, TraesCS1D02G021100 and TraesCS1D02G021200) were localized within intervals of SNP markers associated with biocontrol efficacy Zt - ZtCr (Table 3). TraesCS1D02G021000 and TraesCS1D02G021200 were predicted to encode proteins with sequence similarity to Pik-2-like disease resistance proteins (Table 3). Other proteins present in the region were predicted to be involved in transcriptional regulation, mechanosensitive ion channels and oxidoreductase activities (Table 3).

## 4 Discussion

Exploration of genetic variability through breeding is one of the main ways to improve yield, quality, disease resistance and abiotic stress tolerance in agricultural plants. Likewise, novel traits of interest such as interactions with beneficial microorganisms, interactions with other plants and microbiome modulation can also benefit from a deeper understanding of genetic variation and genotype-genotype interactions. In this study, we explored the natural variation present in 202 winter wheat genotypes for STB disease resistance and variation in wheat genotypes affecting the biocontrol efficacy of *C. rosea* in controlling STB.

Using a large and diverse panel of winter wheat genotypes primarily from the Scandinavian countries, we found significant variation among wheat genotypes for both disease resistance and biocontrol efficacy of *C. rosea*. This panel of winter wheat genotypes has previously been used to explore genetic variation for several diseases such as powdery mildew, fusarium head blight and STB (Odilbekov *et al*. 2019; Alemu *et al*. 2021; Zakieh *et al*. 2021). Our results for STB resistance are in agreement with data reported in Odilbekov *et al*. (2019).

*Clonostachys rosea* is primarily considered as a soil-borne and rhizosphere-associated fungus that has shown biocontrol properties against a multitude of diseases (Jensen *et al*. 2021). However, *C. rosea* is reported to act as a BCA against several diseases in the phyllosphere caused by pathogens such as *F. graminearum*, *Puccinia triticana*, *P hordei*, *P. coronata* f. sp. *avenacea* and *B. sorokiniana* (Xue *et al*. 2009; Jensen *et al*. 2016a; Wilson *et al*. 2020; Jensen *et al*. 2021). In this study, we show that application of *C. rosea* strain IK726 to wheat leaves efficiently protects against STB disease caused by *Z. tritici* under controlled conditions. Biocontrol of STB by *C. rosea* strain IK726 is previously reported from multi-year field trials in Denmark, where *C. rosea* IK726 alone and in combination with other BCAs showed significant reduction in STB compared to untreated control (Jensen *et al*. 2019).

The importance of plant genotypes in establishment and efficient biocontrol and biostimulation from beneficial microorganisms is suggested before (Stenberg *et al*. 2015; Jensen *et al*. 2016b). However, knowledge about the extent and strength of this phenomenon is limited as only a small number of studies, with low numbers of plant genotypes tested, confirm plant host genotype-specific interactions with BCAs and biostimulants (Tucci *et al*. 2011; Moraga-Suazo *et al*. 2016; Prashar & Vandenberg 2017; Schmidt *et al*. 2020). Here, we use 183 wheat genotypes to show that the efficacy of *C. rosea* strain IK726 in controlling STB is quantitatively modulated by the plant genotype. The moderate positive correlation between disease resistance and biocontrol efficacy shows that susceptible plant genotypes typically benefit more from *C. rosea* application. However, the fact that this correlation is not strong shows that there is an additional level of genetic predisposition in the wheat material to benefit from the BCA treatment.

Large scale genetic studies in wheat have found several quantitative trait loci (QTLs) throughout the wheat genome associated with STB disease resistance (Brown *et al*. 2015). Odilbekov *et al*. (2019) found QTLs associated with STB on chromosomes 1A, 1B, 2B, 3A and 5A. In this study, we identified two significant marker trait associations on chromosome 2B and 3B, which is also reported previously on these chromosomes (Brown *et al*. 2015; Odilbekov *et al*. 2019; Alemu *et al*. 2021). Various QTLs on these chromosomes are suggested to contribute to disease resistance specifically at seedling stage and at adult plant stage, reflecting the complexity and variety of putative resistance genes linked to STB (Brown *et al*. 2015; Odilbekov *et al*. 2019). In these regions, several genes predicted to have a role in plant defenses were also localized. The TraesCS3B02G307000 gene is predicted to encode for plant homeodomain (PHD) Zinc finger-type protein known to transcriptionally regulate plant defense genes (Korfhage *et al*. 1994). TraesCS3B02G307300 is a gene predicted to encode for a VQ motif-containing protein, which are known to regulate various developmental process including responses to biotic and abiotic stress generally in all plants (Jing & Lin 2015) and in wheat (Zhang *et al*. 2022). TraesCS3B02G307100 is a predicted dynamin-like GTPase similar to members of dynamin superfamily that are involved in budding of transport vesicles, division of organelles, cytokinesis and pathogen resistance by mediating vesicle trafficking (Praefcke & McMahon 2004). The presence of these resistance sources in Nordic wheat germplasm reflects a genetic potential that can be utilized in breeding to improve STB resistance in wheat.

We further identify significant associations of SNP markers to *C. rosea* biocontrol on chromosome 1B and 6D and biocontrol efficacy on chromosomes 1D and 6B, which are distinctive from marker-trait associations found for disease resistance. These results indicate that the QTLs contributing to STB disease resistance and biocontrol are located in different genomic regions, which suggest that it is possible to breed wheat genotypes that combine high STB disease resistance with high BCA compatibility. SNP markers that exhibit segregation with distinct disease resistance and biocontrol related traits can aid in plant breeding by enabling the simultaneous and more efficient selection of multiple QTLs, offering a cost-effective approach. The underlying mechanisms for this plant genotype-dependent effect on biocontrol efficacy is currently unknown. We identified two genes (TraesCS1D02G021000 and TraesCS1D02G021200) predicted to encode Pik-2 like disease resistance proteins, which are known R proteins inducing hypersensitive response in plants to restrict pathogen growth (Ashikawa *et al*. 2008). The presence of Pik-2-like disease resistance protein genes in regions segregating with biocontrol efficacy may suggest differential ability of wheat genotypes to recognize microbe-associated molecular patterns (MAMPs) or microbial effectors and subsequently in their ability to induce pattern-triggered immunity (PTI) or effector-triggered immunity (ETI) (Jones & Dangl 2006; Köhl *et al*. 2019; Jensen *et al*. 2021). This is in line with results from Moraga-Suazo *et al*. (2016) where *C. rosea*-mediated biocontrol of the pitch canker pathogen *F. circinatum* differs between *P. radiata* genotypes, which in turn is related with the ability of *C. rosea* to activate ISR. In fact, *C. rosea* can trigger defense gene expression in both wheat and tomato (Mouekouba *et al*. 2014; Wang *et al*. 2019; Kamou *et al*. 2020), which subsequently may trigger ISR as shown in wheat (Roberti *et al*. 2008) and tobacco (Lahoz *et al*. 2004). We further identified a gene (TraesCS1D02G020800) involved in mechanosensitive channel protein that provide protection against hypoosmotic shock (Pivetti *et al*. 2003) and a gene (TraesCS1D02G020900) associated with a Myb-like protein involved in transcriptional regulation by DNA binding (Klempnauer & Sippel 1987). The fact that certain wheat genotypes responded negatively to the application of *C. rosea* in the presence of *Z. tritici* illustrates the delicate balance between BCAs, pathogens and plants at cellular and physiological level (Jensen *et al*. 2021).

## 5 Conclusions

This study highlights the role of plant genotypes in efficient application of BCAs. We show that winter wheat genotypes vary in their ability to benefit from *C. rosea*-mediated biocontrol of STB and that disease resistance and biocontrol efficacy are genetically distinct traits. Breeding plants with a high genetic potential to benefit from the application of beneficial microorganisms can facilitate the transition to agricultural production systems with lower input of chemical fungicides. Future studies to elucidate the underlying mechanisms of this phenomenon can further improve our possibilities to integrate beneficial microorganisms in sustainable agriculture.

## 6 List of abbreviations

AUDPC: Area under disease progress curve; BCA: Biological control agent; GWAS: Genome–wide association study; IPM: Integrated pest management; QTL: Quantitative trait loci; SNP: Single nucleotide polymorphism; STB: Septoria tritici blotch

## 7 Declarations

## Ethics approval and consent to participate

Not applicable

## Consent for publication

Not applicable

## Availability of data and materials

All the data produced in this study are available in supplementary files.

## Authors’ contributions

MK, MD, DFJ and LGB conceived the study. All authors contributed to designing the experiments. SC and MZ performed the experiments. SC performed all the analyses. SC wrote the first draft of the manuscript with assistance from MK. All authors read and approved the manuscript.

## Acknowledgements

The authors thank Tina Henriksson at Lantmännen AB for providing advice on experimental design and selection of wheat material. The authors also thank Johannes Forkman for statistical input and Edoardo Piombo for input on steps for candidate genes identification.

## Supplementary

Supp. Figure 1: Phenotypic distribution of scaled relative area under disease progress curve (rAUDPC) of wheat genotypes in (A) treatment Zt (*Z. tritici* alone) and (B) treatment ZtCr (*Z. tritici* along with *C. rosea*). Points represent the model estimated means and error bars represent 95 % confidence intervals for each genotype. Points with different shapes and color represent the country of origin of wheat genotypes.

Supp. Figure 2: Allelic level comparison of SNP markers significantly associated with scaled relative area under disease progress curve (rAUDPC) of wheat genotypes in treatment Zt (*Z. tritici* alone), treatment ZtCr (*Z. tritici* along with *C. rosea*) and biocontrol efficacy (Zt – ZtCr). Panel A-F display significant markers with their location, their alleles and distribution of wheat genotypes for the associated trait. Alleles not sharing the same letter are significantly different at *P* < 0.05. Numbers at the bottom of the panel indicate number of genotypes, and black diamonds represent mean estimate of the group.

Supp. Table 1: List of 202 winter wheat genotypes from NordGen gene bank with information on country of origin, year of origin and pedigree.

Supp. Table 2: Single nucleotide polymorphism (SNP) marker information for 173 winter wheat genotypes used in the GWAS analyses.

Supp. Table 3: Small scale biocontrol efficacy screening experiment model output across treatments and genotypes.

Supp. Table 4: Model output for the treatment Zt (*Z. tritici* only) of 202 winter wheat genotypes.

Supp. Table 5: Model output for the treatment ZtCr (*Z. tritici* along with *C. rosea*) of 183 winter wheat genotypes.

Supp. Table 6: Biocontrol efficacy estimates (Zt - ZtCr) of *C. rosea* in controlling septoria tritici blotch for 183 winter wheat genotypes.

Supp. Table 7: Gene content of wheat genomic regions segregating with septoria tritici blotch disease resistance and biocontrol in ± 1.6 cM interval as per Alemu et. al. (2021)

