## Supplementary figures and images for "Plant genotype-specific modulation of *Clonostachys rosea*-mediated biocontrol of septoria tritici blotch disease on wheat"

### Supp Fig 1

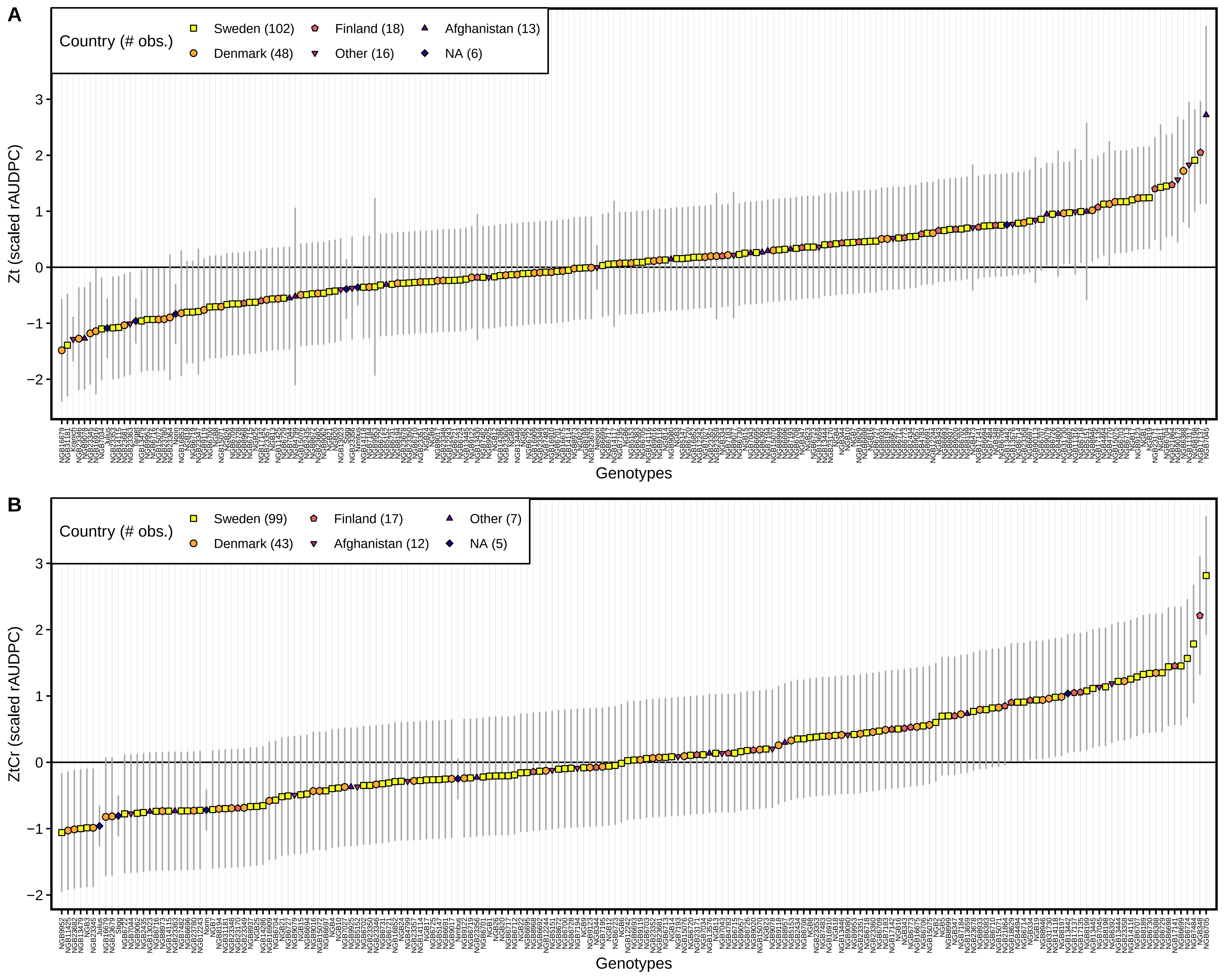

### Supp Fig 2

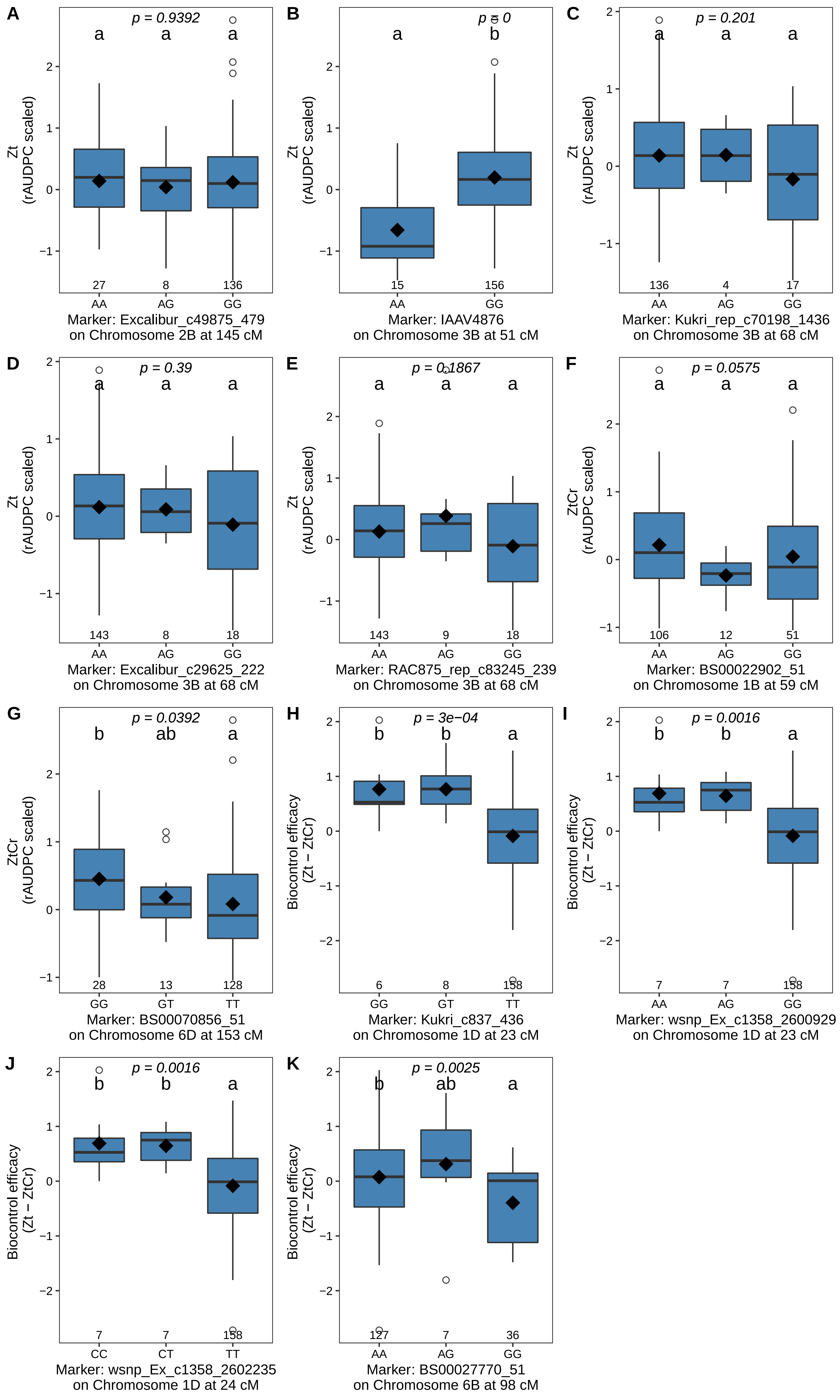
